# Neutralization of SARS-CoV-2 lineage B.1.1.7 pseudovirus by BNT162b2 vaccine-elicited human sera

**DOI:** 10.1101/2021.01.18.426984

**Authors:** Alexander Muik, Ann-Kathrin Wallisch, Bianca Sänger, Kena A. Swanson, Julia Mühl, Wei Chen, Hui Cai, Ritu Sarkar, Özlem Türeci, Philip R. Dormitzer, Ugur Sahin

**Affiliations:** BioNTech, An der Goldgrube 12, 55131 Mainz, Germany; Pfizer, 401 N Middletown Rd., Pearl River, NY 10960, U.S.A.; TRON gGmbH – Translational Oncology at the University Medical Center of the Johannes Gutenberg, University Freiligrathstraße 12, 55131 Mainz, Germany

## Abstract

Recently, a new SARS-CoV-2 lineage called B.1.1.7 has emerged in the United Kingdom that was reported to spread more efficiently than other strains. This variant has an unusually large number of mutations with 10 amino acid changes in the spike protein, raising concerns that its recognition by neutralizing antibodies may be affected. Here, we investigated SARS-CoV-2-S pseudoviruses bearing either the Wuhan reference strain or the B.1.1.7 lineage spike protein with sera of 16 participants in a previously reported trial with the mRNA-based COVID-19 vaccine BNT162b2. The immune sera had equivalent neutralizing titers to both variants. These data, together with the combined immunity involving humoral and cellular effectors induced by this vaccine, make it unlikely that the B.1.1.7 lineage will escape BNT162b2-mediated protection.

## Main Text

In a pivotal phase 3 trial conducted in the United States, Argentina, Brazil, South Africa, Germany, and Turkey, the BioNTech-Pfizer mRNA vaccine BNT162b2 was 95% effective in preventing COVID-19 through the data cut-off date of November 14, 2020 (*1*). In September 2020, the SARS-CoV-2 lineage B.1.1.7 was detected in the United Kingdom (*2*), and it subsequently increased in prevalence, showed enhanced transmissibility, and spread to other countries and continents (*3*). B.1.1.7 has a series of mutations in its spike (S) protein: ΔH69/V70, ΔY144, N501Y, A570D, D614G, P681H, T716I, S982A, and D1118H. One of these mutations, N501Y, was of particular concern because it is located in the receptor binding site, the spike with this mutation binds more tightly to its cellular receptor, ACE-2 (*4*); and virus with this mutation has an increased host range that includes mice (*5*). BNT162b2-immune sera neutralized SARS-CoV-2 (USA/WA-1/2020 background strain) with an introduced N501Y mutation as efficiently as SARS-CoV-2 without the mutation (*6*). Further, 19 pseudoviruses, each bearing a SARS-CoV-2 S with a different mutation found in circulating strains, were also neutralized as efficiently as non-mutant SARS-CoV-2 S bearing pseudoviruses by BNT162b2-immune sera (*7*). Nevertheless, the question remained whether a virus with the full set of mutations in the lineage B.1.1.7 spike would be neutralized efficiently by BNT162b2-immune sera.

To answer this question, we generated VSV-SARS-CoV-2-S pseudoviruses bearing the Wuhan reference strain or the lineage B.1.1.7 spike protein (Fig S1). Sera of 16 participants in the previously reported German phase 1/2 trial (*7*), drawn from eight younger (18-55 yrs) and eight older adults (56-85 yrs) at 21 days after the booster immunization with 30 μg BNT162b2 (Fig S2), were tested for neutralization of SARS-CoV-2 Wuhan and lineage B.1.1.7 spike-pseudotyped VSV by a 50% neutralization assay (pVNT_50_). The ratio of the 50% neutralization geometric mean titer (GMT) of the sera against the SARS-CoV-2 lineage B.1.1.7 spike-pseudotyped VSV to that against the Wuhan reference spike-pseudotyped VSV was 0.79, indicating no biologically significant difference in neutralization activity against the two pseudoviruses (Fig. 1, Fig. S3, Table S1).

**Fig. 1.**
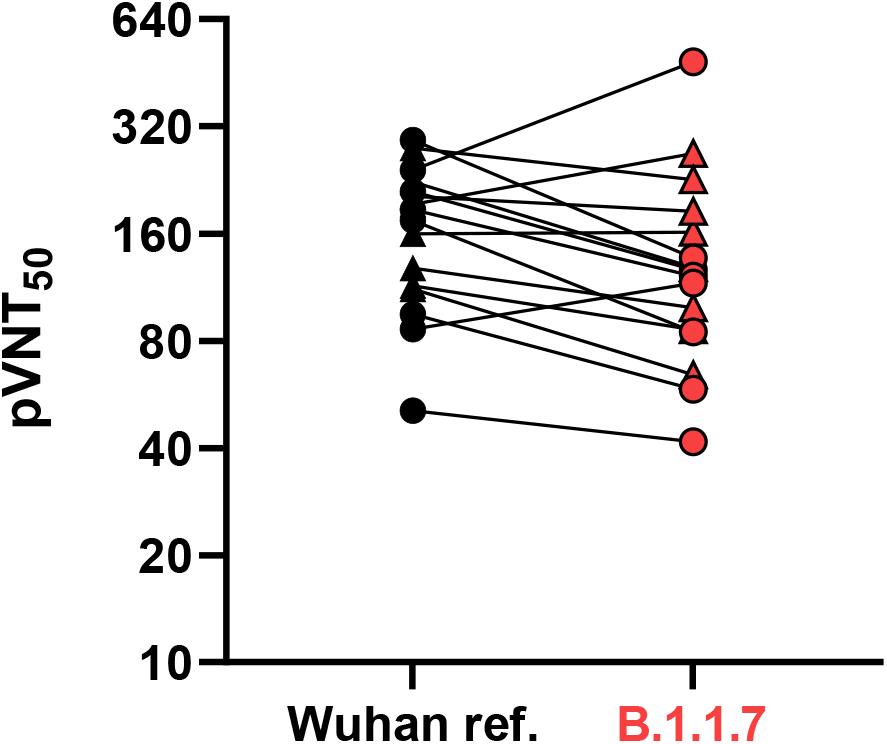
50% pseudovirus neutralization titers of 16 sera from BNT162b2 vaccine recipients against VSV-SARS-CoV-2-S pseudovirus bearing the Wuhan or lineage B.1.1.7 spike protein. N=8 representative sera each from younger adults (aged 18 to 55 yrs; indicated by triangles) and older adults (aged 56 to 85 yrs; indicated by circles) drawn at day 43 (21 days after dose 2) were tested.

The preserved neutralization of pseudoviruses bearing the B.1.1.7 spike by BNT162b2-immune sera makes it very unlikely that the UK variant viruses will escape BNT162b2-mediated protection. A potential limitation of the work may be the use of a non-replicating pseudovirus system. However, previous reports have shown good concordance between pseudotype neutralization and SARS-CoV-2 neutralization assays (*8, 9*). Additional experiments will confirm efficient neutralization of B.1.1.7 lineage clinical isolates. The ongoing evolution of SARS-CoV-2 necessitates continuous monitoring of the significance of changes for maintained protection by currently authorized vaccines. Unlike for influenza vaccines, the reduction in neutralization that might indicate the need for a strain change has not been established for COVID-19 vaccines. Given the polyepitopic immune response with concurrent activation of neutralizing antibodies and T cells, and thus multiple potential mediators of protection elicited by BNT162b2, it is possible that vaccine efficacy could be preserved, even with substantial losses of neutralization by vaccine-elicited sera. Nevertheless, preserved neutralization is reassuring, and preparation for potential COVID-19 vaccine strain change is prudent. Such an adaptation of the vaccine to a new virus strain would be facilitated by the flexibility of mRNA-based vaccine technology.

## Acknowledgments

Supported by BioNTech and Pfizer. We thank the BioNTech German clinical trial (NCT04380701, EudraCT: 2020-001038-36) participants, from whom the post-immunization human sera were obtained. We thank the many colleagues at BioNTech and Pfizer who developed and produced the BNT162b2 vaccine candidate.

## Author contributions

U.S., Ö.T., A.M. and P.R.D. conceived and conceptualized the work. K.A.S. and A.M planned and supervised experiments. A.M., A.W., J.M., B.S., H.C, W.C. and R.S. performed experiments. A.M., H.C, and K.A.S. analyzed data. U.S., Ö.T., A.M., P.R.D. and K.A.S. interpreted data and wrote the manuscript. All authors supported the review of the manuscript.

## Data and materials availability

The data that support the findings of this study are available from the corresponding author upon reasonable request.

