## Supplementary Material for "Neutralization of SARS-CoV-2 lineage B.1.1.7 pseudovirus by BNT162b2 vaccine-elicited human sera"

### Table of Contents


#### Materials and Methods

- VSV-SARS-CoV-2 S variant pseudovirus generation
- Serum specimens and neutralization assay

#### Figures

- Fig. S1. Schematic illustration of the production of VSV pseudoviruses bearing SARS-CoV-2 S protein.
- Fig. S2. Scheme of the BNT162b2 vaccination and serum sampling.
- Fig. S3. Plot of the ratio of pVNT50 between SARS-CoV-2 lineage B.1.1.7 and Wuhan reference strain spike-pseudotyped VSV.
- Fig. S4. Titration of SARS-CoV-2 Wuhan reference strain and lineage B.1.1.7 spike-pseudotyped VSV on Vero 76 cells using GFP-infected cells as read-out

#### Tables

- Table S1. pVNT50 values of 16 BNT162b2 post-immunization sera against SARS-CoV-2 Wuhan reference strain spike-pseudotype and lineage B.1.1.7 spike-pseudotyped VSV.
- Table S2. Titers of SARS-CoV-2 Wuhan reference strain and lineage B.1.1.7 spike-pseudotyped VSV in transducing units (TU) per mL.

#### References

### Materials and Methods

VSV-SARS-CoV-2 S variant pseudovirus generation

A recombinant replication-deficient vesicular stomatitis virus (VSV) vector that encodes green fluorescent protein (GFP) and luciferase instead of the VSV-glycoprotein (VSV-G) was pseudotyped with SARS-CoV-2 spike (S) derived from either the Wuhan reference strain (NCBI Ref: 43740568) or the variant of concern (VOC)-202012/01 (also known as SARS-CoV-2 lineage B.1.1.7) according to published pseudotyping protocols (*10*). In brief, HEK293T/17 monolayers transfected to express SARS-CoV-2 S were inoculated with VSV-G complemented VSVΔG vector. After incubation for 1 h at 37 °C, the inoculum was removed. Cells were washed with PBS before medium supplemented with anti-VSV-G antibody (clone 8G5F11, Kerafast Inc.) was added to neutralize residual VSV-G complemented input virus. VSV-SARS-CoV-2-S pseudotype-containing medium was harvested 20 h after inoculation, passed through a 0.2 µm filter and stored at -80 °C. Prior to use in the neutralization test, the pseudovirus batches were titrated on Vero 76 cells, and the percent infected cells determined by flow cytometry (Fig. S4). Individual titers were calculated in transducing units (TU) per mL. Production of the VSV-SARS-CoV-2-S pseudoviruses bearing the Wuhan reference strain or lineage B.1.1.7 strain spike protein yielded similar titers (Table S2).

Serum specimens and neutralization assay

The immunization was performed on days 1 and 22, and serum was collected at both time points and additionally on day 43 (3 weeks after dose 2). For measuring neutralization titers, each serum collected at day 43 was 2-fold serially diluted in culture medium with the first dilution of 1:20 (dilution range of 1:20 to 1:2560). VSV-SARS-CoV-2-S particles were diluted in culture medium to obtain 1,000 TU in the assay. Serum dilutions were mixed 1:1 with pseudovirus for 30 minutes at room temperature prior to addition to Vero 76 cell monolayers in 96-well plates and incubation at 37 °C for 24 hours. Supernatants were removed, and the cells were lysed with luciferase reagent (Promega). Luminescence was recorded, and neutralization titers were calculated in GraphPad Prism version 9 by generating a 4-parameter logistical (4PL) fit of the percent neutralization at each serial serum dilution. The 50% pseudovirus neutralisation titre (pVNT50) was reported as the interpolated reciprocal of the dilution yielding a 50% reduction in luminescence. A table of the neutralization titers is provided (Table S2). The ratio for each serum of the pVNT50 against SARS-CoV-2 lineage B.1.1.7 and the Wuhan reference strain spike-pseudotyped VSV is plotted in Fig. S3.

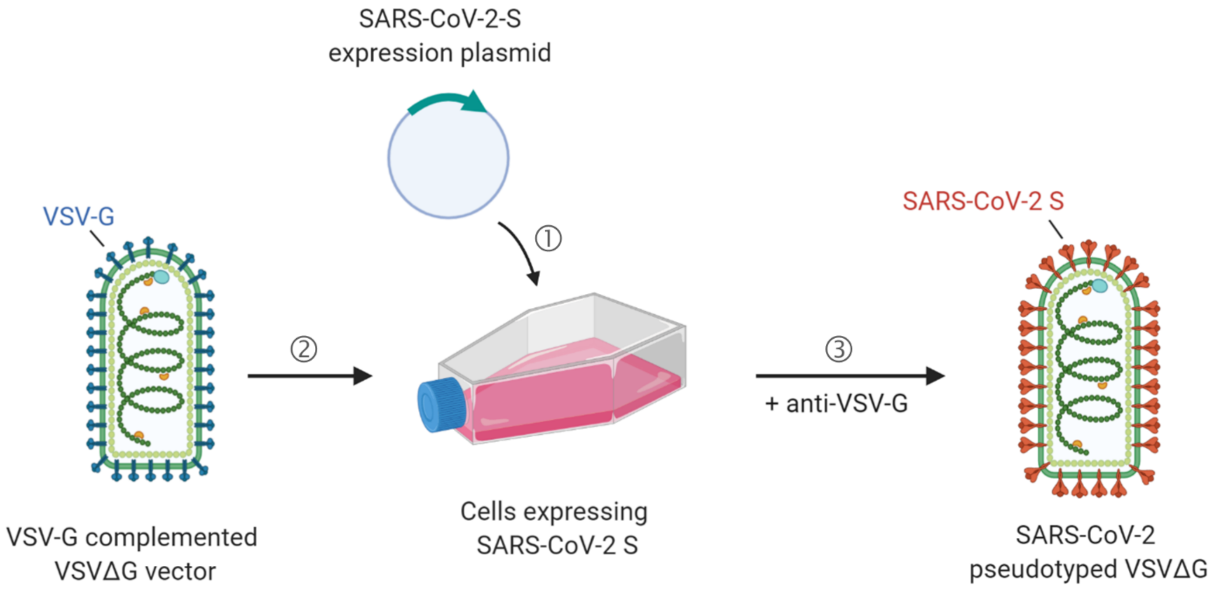


**Fig. S1. Schematic illustration of the production of VSV pseudoviruses bearing SARS-CoV-2 S protein.** (1) Transfection of SARS-CoV-2-S expression plasmid into HEK293/T17 cells. (2) Infection of SARS-CoV-2 S expressing cells with VSV-G complemented input virus lacking the VSV-G in its genome (VSVΔG) and encoding for reporter genes. (3) Neutralization of residual VSV-G complemented input virus by addition of anti-VSV-G antibody yields SARS-CoV-2 S pseudotyped VSVΔG as a surrogate for live SARS-CoV-2.

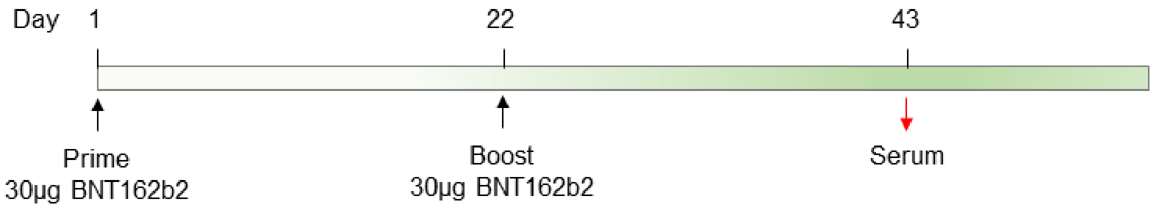


**Fig. S2. Scheme of the BNT162b2 vaccination and serum sampling.**

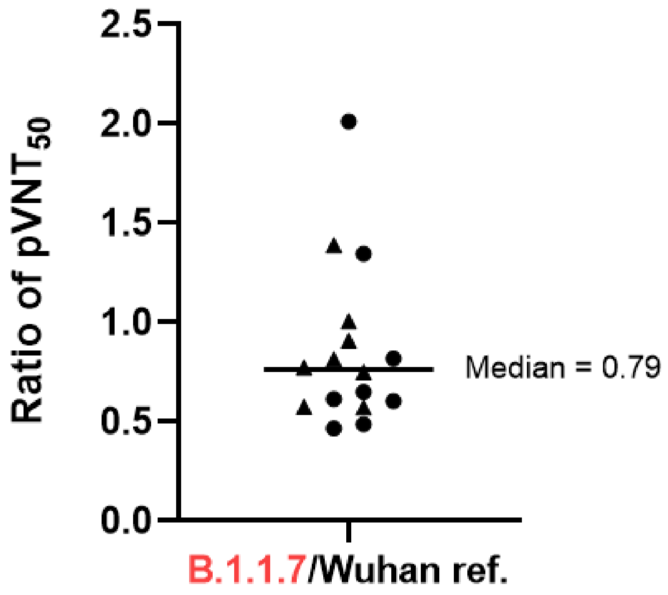


**Fig. S3. Plot of the ratio of pVNT50 between SARS-CoV-2 lineage B.1.1.7 and Wuhan reference strain spike-pseudotyped VSV.** Triangles represent sera from younger adults (aged 18 to 55 yrs), and circles represent sera from older adults (aged 56 to 85 yrs). The sera were drawn on day 43 (21 days after dose 2).

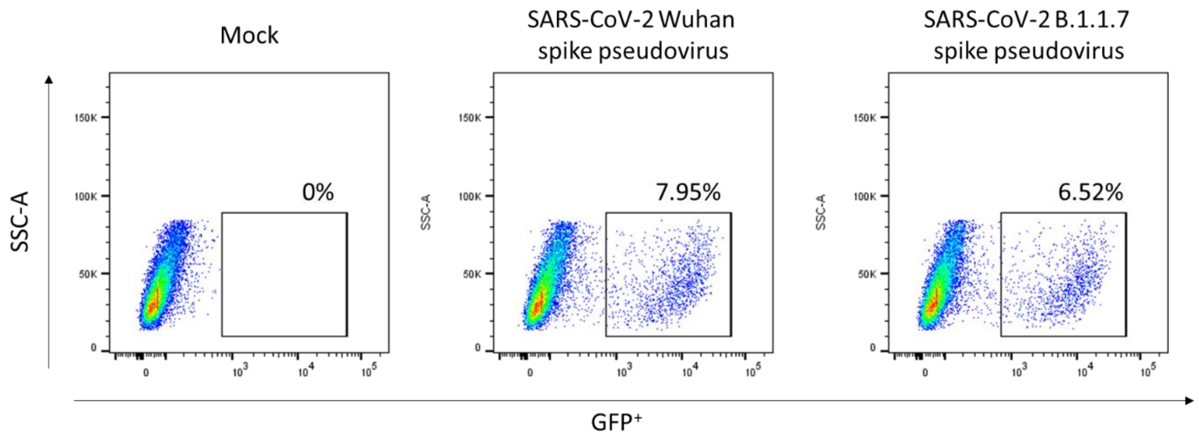


**Fig. S4. Titration of SARS-CoV-2 Wuhan reference strain and lineage B.1.1.7 spike-pseudotyped VSV on Vero 76 cells using GFP-infected cells as read-out.**

**Table S1. pVNT50 values of 16 BNT162b2 post-immunization sera against SARS-CoV-2 Wuhan reference strain spike-pseudotype and lineage B.1.1.7 spike-pseudotyped VSV.**

| Serum ID | $pVNT_{50}$ | | $pVNT_{50}$ ratio |
|---|---|---|---|
| | Wuhan ref. | B.1.1.7 | (B.1.1.7/Wuhan ref.) |
| 1 | 160 | 161.2 | 1.01 |
| 2 | 114.1 | 85.8 | 0.75 |
| 3 | 223.2 | 128.6 | 0.58 |
| 4 | 193 | 268.4 | 1.39 |
| 5 | 111.9 | 64.3 | 0.57 |
| 6 | 128 | 99.1 | 0.77 |
| 7 | 278.1 | 226.8 | 0.82 |
| 8 | 203.6 | 185 | 0.91 |
| 9 | 94.9 | 58.4 | 0.62 |
| 10 | 209.7 | 126.8 | 0.60 |
| 11 | 50.8 | 41.7 | 0.82 |
| 12 | 241.3 | 486.1 | 2.01 |
| 13 | 174 | 84.8 | 0.49 |
| 14 | 292.5 | 136.7 | 0.47 |
| 15 | 186.7 | 121.6 | 0.65 |
| 16 | 86.3 | 116.2 | 1.35 |

**Table S2. Titers of SARS-CoV-2 Wuhan reference strain and lineage B.1.1.7 spike-pseudotyped VSV in transducing units (TU) per mL.**

| VSV pseudovirus bearing | Titer [TU/mL] |
|---|---|
| Wuhan strain SARS-CoV-2 S | $1.59x10^5$ |
| Lineage B.1.1.7 SARS-CoV-2 S | $1.30\ x10^5$ |
